# Learning a CoNCISE language for small-molecule binding

**DOI:** 10.1101/2025.01.08.632039

**Authors:** Mert Erden, Kapil Devkota, Lia Varghese, Lenore Cowen, Rohit Singh

**Author notes:** These authors contributed equally (Mert Erden, Kapil Devkota).

## Abstract

Rapid advances in deep learning have improved <u>in silico</u> methods for drug-target interaction (DTI) prediction. However, current methods do not scale to the massive catalogs that list millions or billions of commercially-available small molecules. Here, we introduce CoNCISE, a method that accelerates drug-target interaction (DTI) prediction by 2-3 orders of magnitude while maintaining high accuracy. CoNCISE uses a novel vector-quantized codebook approach and a residual-learning based training of hierarchical codes. Strikingly, we find that much of binding-specificity information in the small molecule space can be compressed into just 15 bits of information per compound, characterizing all small molecules into 32,768 hierarchically-organized binding categories. Our DTI architecture, which combines these compact ligand representations with fixed-length protein embeddings in a cross-attention framework, achieves state-of-the-art prediction accuracy at unprecedented speed. We demonstrate CoNCISE’s practical utility by indexing 6.4 billion ligands in the Enamine dataset, enabling researchers to query vast chemical libraries against a protein target in seconds. A “CoNCISE + docking” pipeline screened Enamine to propose strong binders (predicted *K*_*D*_ ≈ 10-20 *µ*M) of three difficult-to-drug targets, each within two hours. CoNCISE’s advance could democratize access to largescale computational drug discovery, potentially enabling rapid identification of promising molecules for therapeutic targets and cellular perturbations.

## 1 Introduction

Small molecules are powerful therapeutic and basic-science reagents, offering shelf stability, oral bioavailability, and precise temporal control in cell perturbation experiments. While a successful drug eventually needs to satisfy diverse criteria for safety and efficacy [23], an essential first requirement for a small molecule drug is binding: given a protein target, we need to find molecules that bind strongly to it. This is termed the <u>drug-target interaction prediction</u> (DTI) problem. The space of possible small molecules is so vast, however, that even for proteins that have strong-binding ligands somewhere in chemical space, finding them is a massive challenge. Some researchers take a *generative* approach to construct novel candidate binders [19], but here we focus on the complementary *discriminative* approach of screening existing molecule libraries.

Unlike peptides, small molecule synthesis requires a customized protocol for each compound, making de novo synthesis inaccessible to many researchers. The alternative is to screen libraries of existing or readily-synthesizable molecules to identify candidates for experimental evaluation [17,40]. Massive catalogs like ZINC [18] and Enamine [13] now list millions and billions, respectively, of commercially-available small molecules. However, accurately and efficiently screening these massive libraries even for a single protein target is an open problem.

Existing computational approaches to DTI prediction are too slow for screening massive molecular databases. Traditional approaches rely on structural docking to estimate binding affinities and identify optimal conformations [4,10], but this is computationally very expensive. Recent deep learning-based docking techniques [5] leveraging predicted structures [20,42,21] are faster but still require about one minute per DTI. An alternative, much faster approach has been enabled by the use of sequence-based representations, namely, protein language models (PLMs) and molecular fingerprints. These represent targets and drugs in a high-dimensional space where neural networks learn implicit biophysical patterns of interaction. PLMs like Bepler-Berger [2,3], ESM [30,22] and ProtBert [9] encode proteins in embeddings that implicitly capture structural information. Our recent PLM-based method ConPLex [32], which encoded drugs using Morgan fingerprints [29] of SMILES strings [28], accurately screens DTIs by mapping proteins and drugs into a shared embedding space. While ConPLex is much faster than previous approaches (0.01-0.001 secs/DTI), screening all of ZINC or Enamine would still take days or weeks for a single protein target.

Understanding the underlying structure of chemical space could dramatically accelerate drug screening. For instance, ConPLex learned to co-embed proteins and drugs into a 1024-dimensional space, where cosine distances predict binding affinity. However, this space is still vast and, with only ∼ 36, 000 DTIs used to train ConPLex, sparsely occupied. We wondered if it might be possible to learn a much more compact co-embedding that is as accurate and interpretable as ConPLex’s, but additionally provides an efficient, informative way to organize the small molecule space.

We hypothesized that recent advances in interpretability of neural networks [36] could offer both unprecedented insights into small-molecule space and dramatic acceleration of DTI prediction. **Our key conceptual advance is to create a binding-informed hierarchical clustering of the entire small-molecule space**, by introducing the use of vector-quantized (VQ) “codebooks” for small molecule representation (**Figure 1A-C**). Quantization has proven powerful in domains ranging from language models to protein structure analysis [38,33], including tokenization in Foldseek [39] and the PLMs ESM-3 [16] and SaProt [35].

**Fig. 1:**
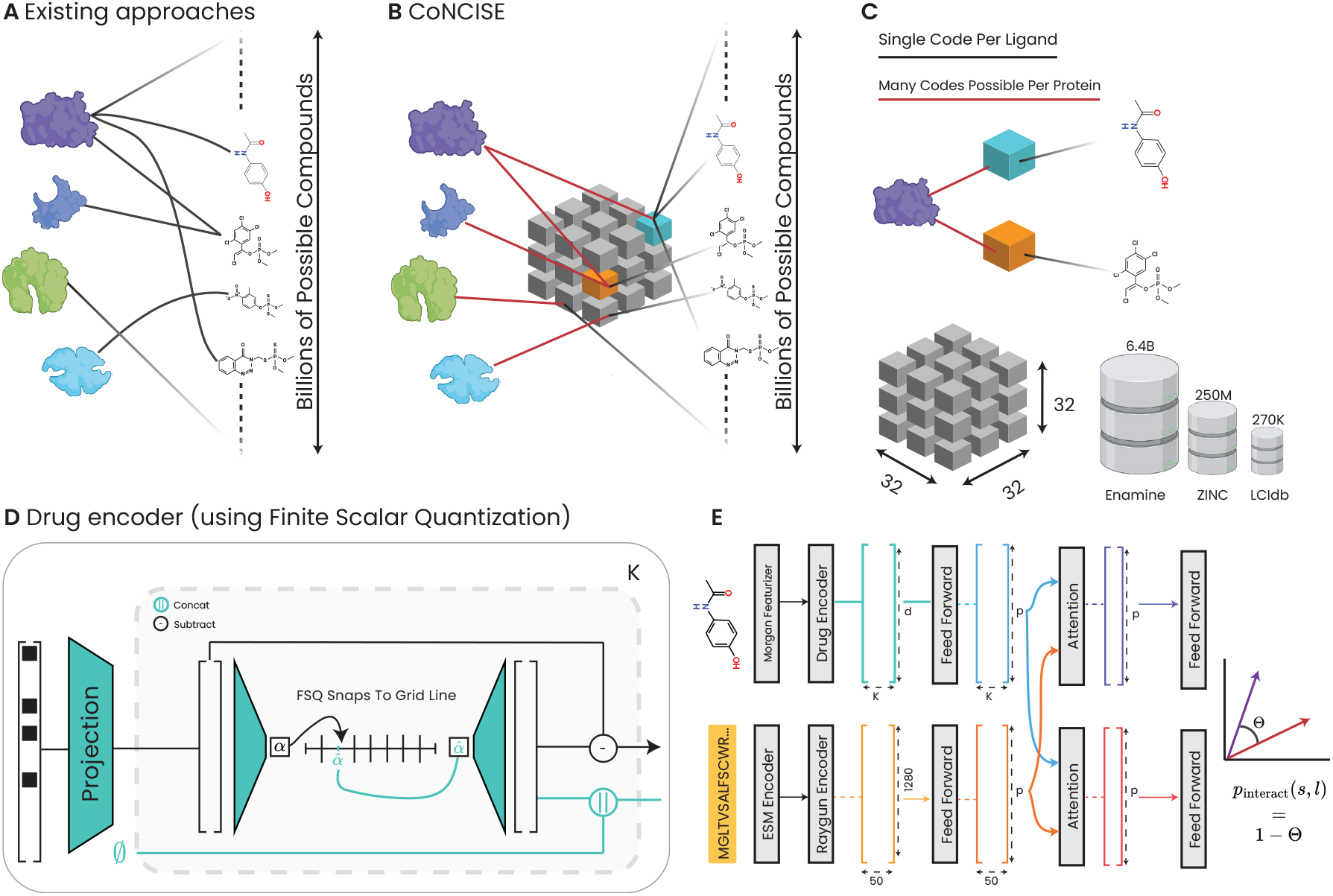
Overview. A) Conventional DTI search requires evaluating all possible protein-drug pairs. B) CoNCISE instead compresses drug space into a finite codebook of 32^3^ representations, enabling efficient binding prediction through codes. C) Each target’s binding profile becomes a concise list of codes, enabling constant-time database queries regardless of size. D) CoNCISE’s novel drug encoder takes binary fingerprints, down-projects them, and extracts *K* many discrete points that represent the ligand. Discretization to the compressed codebook space happens by snapping the low-dimensional representation onto grid lines. Each block produces a quantized representation, and a residual that is propagated (*K* = 3 levels, grid length = 32). E) The complete CoNCISE architecture.

We introduce CoNCISE (Compact Novel Codebook Interaction Sequence Embeddings), which implements this codebook approach through a novel residual-learning framework. CoNCISE quantizes the vast small molecule space to 32, 768 (=2^15^) representations, expressed through a hierarchical 3-tier “ligand code”: *a*.*b*.*c*, where *a, b*, and *c* are limited to 32 (=2^5^) values each (**Figure 1C**). This allows us to represent any ligand with 15 bits of information, offering many-fold speed and space advantage over the traditional Morgan fingerprint’s 2^2048^ representation space. In particular, with compressed representations we can store SMILES strings of all of Enamine in 168GB, and we can index them by CoNCISE codes that require 16GB of space (whereas indexing via Morgan fingerprints would need 361GB). Crucially, codes are optimized for DTI predictability rather than being derived from unsupervised clustering of drug features, and can be computed in constant time, as our neural network directly outputs the code given an input SMILES string.

An additional innovation in CoNCISE is integrating the ligand representation with Raygun, a recently-described fixed-length protein embedding capturing overall protein structure [7]. Our DTI architecture uses the protein’s sequence and the drug’s SMILES descriptor as input, converts them to the Raygun embedding and Morgan fingerprints, respectively, then applies the codebook architecture on the drug, and finally relates the drug code and Raygun representation in a cross-attention framework.

CoNCISE achieves state-of-the-art prediction accuracy in sequence-based DTI prediction at unprecedented speed. Notably, an understanding of the drug space and protein binding profiles <u>emerges</u> from the model: i) Tanimoto similarity increases along the hierarchy, and ii) when evaluated on the human proteome, a wide range of binding promiscuity is observed– the median protein binds 144 codes strongly, but a small set of proteins has no strong binders while another small set has >1,000 binding codes. To demonstrate CoNCISE’s efficiency and utility, we indexed 6.4B ligands in Enamine. This opens the possibility for researchers to rapidly query vast chemical libraries against protein targets, potentially democratizing access to large-scale computational drug discovery. As a proof of concept, we demonstrate a CoNCISE →DiffDock→SwissDock virtual screening pipeline that retrieves strong predicted binders of KRAS, CXCR4, and P53 from Enamine.

CoNCISE represents a significant advance in our ability to efficiently explore and understand chemical space in the context of drug discovery. It could enable a world where a researcher with a mouse model of a rare disease can query their protein target against massive compound libraries, identify promising molecules within seconds, and order them for testing– a capability that has been infeasible until now.

## 2 Methods

CoNCISE predicts drug-target interactions using two inputs: a protein sequence *S* and a drug’s SMILES representation *L*. While only the protein’s sequence is needed as input, we operate on its PLM-derived (here, ESM-2 [22]) embedding that implicitly captures structural information. The key innovation of CoNCISE, distinguishing it from previous approaches like ConPLex [32], Komet [14] or EnzPred [12], is its requirement that all interaction predictions use an intermediate *discrete* drug representation. This discretization is critical to scalability: once a database has been pre-processed, querying any protein against it requires constant time. It also is important for interpretability and generalizability, i.e., extending from ∼ 500, 000 training examples to billions of ligands. The key design considerations in the CoNCISE formulation then are the discretization scheme, the choice of drug and protein representations, and the overall architecture computing the interaction probability. We start by providing some general background on discretization techniques for neural networks and then detail our specific approach.

### Background on quantized representation learning

Neural networks typically operate on continuous-valued spaces, where features can take on arbitrary values in ℝ^*n*^. While this allows flexibility, it poses challenges for learning compact, interpretable representations. To address this, autoencoders and variational autoencoders learn compact—albeit continuous—latent spaces, but these can still be difficult to index or interpret efficiently. Vector quantization, implemented in architectures like VQ-VAE [38], offers a solution by constraining the latent space to a discrete set of possible values (“codes”). It maps continuous vectors to a finite set of discrete codes through a nearest-neighbor lookup in a learned codebook. This enhances interpretability and sparsity and has proven especially powerful in representing protein structure as a sequence, as demonstrated in methods like Foldseek and SaProt [39,35]. However, traditional VQ-VAEs can be difficult to train and often suffer from “code collapse,” where significant parts of the codebook remain unused.

We instead follow Mentzer et al. [27], who recently proposed a conceptually simple alternative, Finite Scalar Quantization (FSQ): each dimension of the latent space is mapped to a bounded range and then uniformly segmented, leading to *n*-dimensional hypercube whose vertices are the discrete locations. Any point in this space is mapped to its nearest quantization through rounding. While rounding produces no gradient, straight-through estimation enables training. Formally, for each axis *i*, the interval [−1, 1] is divided into *C*_*i*_ uniform regions, yielding 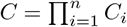 total discrete points (**Appendix 1**). Mentzer et al. document how FSQs are easier to train and do not suffer from code collapse. However, the right codebook size and latent dimensionality is problem-dependent and we next considered the most appropriate settings for CoNCISE.

### Considerations of codebook size

The design of our codebook involves balancing three key factors: the number of distinct protein-drug binding mechanisms that potentially exist, the amount of available training data, and the computational feasibility of model training. While traditional molecular fingerprints like Morgan fingerprints [29] are also codes, their vast space (=2^2048^ codes) is impractical for efficient lookup or learning. To estimate an appropriate size, we performed Fermi calculations focused on the human proteome. Swiss-Prot reports 20,428 human proteins (53,476 including isoforms). While a drug could potentially bind a protein in myriad ways, many drugs likely share similar binding mechanisms against a particular protein, as evidenced by the existence of active sites. Assuming 1-10 such mechanisms per protein, and noting that some mechanisms might be similar across proteins, we estimate between 20,000 and 500,000 distinct binding patterns. This suggests codebook sizes between 2^14^ and 2^19^ would be appropriate. However, with our DTI dataset containing only 111,210 distinct ligands, the sparsity of ligand coverage in the larger codebooks is a concern. We therefore chose 2^15^ (32,768) codes as the largest feasible size. Ablations show this choice performed better than others (**Table 3**).

### Hierarchical codes in CoNCISE

We address the challenge of learning this large codebook through an innovative hierarchical architecture. Inspired by principal component analysis, where each component explains an orthogonal subspace of the data, we layer three FSQ blocks to learn successive one-dimensional latent spaces. In each block, an in- and out-projection layer sandwich a parameter-free FSQ layer mapping to a one-dimensional latent space (**Appendix A**). Each block trains only on the residual error of previous blocks, ensuring it captures complementary information. This residual learning approach naturally induces a tree topology: the first layer makes one of 32 choices for coarse-grained categorization, with the two subsequent layers providing increasingly fine-grained distinctions. Together, these specify a 3-level hierarchical code representation for any ligand. <u>To our knowledge, our innovation of organizing multiple FSQ blocks in a residual, hierarchical format is a novel deep learning contribution that could be broadly useful in learning informative, data-efficient codes</u>. It not only enables efficient, generalizable indexing but also provides an interpretable clustering of the chemical space. Empirical validation confirms this hierarchical organization: molecules sharing partial codes demonstrate greater Tanimoto similarity [1] (**Figure 2**).

**Fig. 2:**
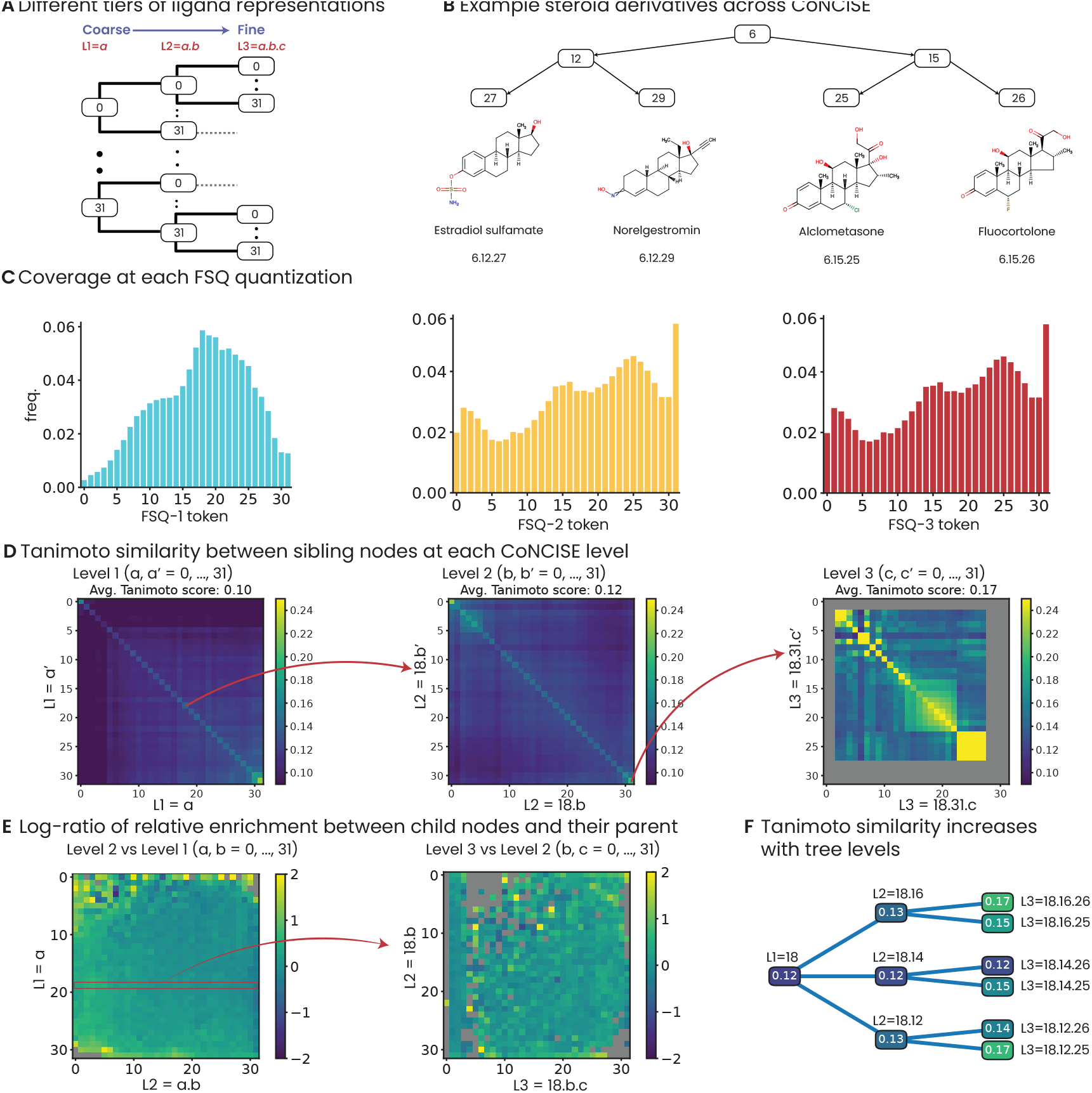
Hierarchical organization of chemical space in CoNCISE. A) The three-tier FSQ hierarchy enables increasingly fine-grained representations. B) Related steroid derivatives cluster together, sharing codes at top levels. C) All FSQ layers show balanced code utilization, avoiding collapse. D) Tanimoto similarity increases between sibling nodes at deeper levels, shown here using representative subtrees (18.* for L2, 18.31.* for L3). Gray indicates insufficient data. E) Child nodes show consistent enrichment over parent nodes in structural similarity. F) Quantitative demonstration of increasing Tanimoto scores along example paths in the hierarchy.

### Drug representation

While SMILES strings are human readable, their variable length and linear nature complicate learning tasks. The field instead typically uses fixed-length fingerprints, like Morgan fingerprints [31], which capture molecular substructures by hashing identical substructures to the same positions in a binary vector. Morgan fingerprints require two parameters: radius (controlling global molecular similarity) and vector length (limiting hash collisions). Following ConPLex [32], we set CoNCISE’s Morgan featurizer to radius 2 and length 2048.

### Protein representation

Protein language models (PLMs), pre-trained on millions of sequences, have emerged as powerful tools for capturing protein structure and function in learned representations [3,22]. Recent methods like ConPLex have shown that these representations can effectively predict drug-target interactions when combined with molecular fingerprints [32]. However, most PLM-based approaches rely on average pooling to obtain fixed-length embeddings, which has two key limitations. First, by aggregating across entire protein, average pooling obscures locations where drug interactions might occur. Second, it attenuates the signal from individual amino acid positions, making it difficult to perform in silico mutagenesis– a mutation’s effect in a 1,000-residue protein might be lost in the averaged representation. To address these limitations, CoNCISE uses Raygun embeddings [7], which encode any protein as 50 contiguous chunks in ℝ^50×1280^. This fixed-length representation preserves local structural information while enabling efficient computation. Crucially, the representation remains sensitive to local changes while capturing global structure.

### 2.1 DTI prediction architecture

The overall CoNCISE architecture consists of a drug encoder module that quantizes the input drug fingerprints into *K* discrete codebooks of size *C* (*K* = 3, *C* = 32 in our final implementation), a protein encoder module that further transforms the Raygun embeddings to enrich for DTI specific information, and the final prediction block, that combines the obtained drug and protein representations to produce a final interaction score. Each of these blocks are described below:

#### Drug Encoder (Figure 1d)

The drug encoder performs quantization through *K* FSQ blocks. Each FSQ block is composed of a parameter-free FSQ layer, which is placed in between two linear projection layers. Let 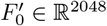 be the drug fingerprint and *F*_0_ ∈ ℝ^*d*^ be the its initial projection to a much smaller dimension *d* << 2048. Additionally, let *FSQ*_*i*_, *i* ∈ {1, …, *K*} be the *K* FSQ blocks. Then, the residuals *F*_*i*_ and the quantizations *Q*_*i*_ are computed as follows:

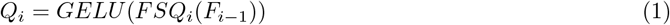

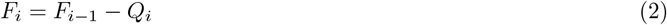

where *GELU* is the non-linear activation employed. Finally, each block’s quantization is concatenated to produce the output embeddings ***Q*** = [*Q*_1_, … *Q*_*K*_]^*T*^ ∈ ℝ^*K*×*d*^.

In addition to the quantized embeddings, the parameter-free FSQ layers internally keep track of the internal quantization state, stored as an integer value, during the forward pass. Together, these *K* integers contain all the information needed to reconstruct the *K* × *d* embeddings (Appendix A.3). Consequently, we use the *K* codebook integer representation of a ligand interchangeably with the drug encoder outputs.

##### Notation

The model we selected for inference has 3 FSQ blocks, each with codebook quantizations of size 32; (32^3^ quantizations in total.) We use the term “level” (*L*) to specify the granularity of codes, corresponding to the level of hierarchy they belong in, e.g. for a ligand, *a*.*b*.*c* is its L3 representation, *a*.*b* is its L2 representation.

#### Quantized Ligand-Protein Attention

We learn relationships between proteins and quantized ligands via cross-attention. Coming out of the drug and Raygun encoders, ligands and proteins have a ***Q*** ∈ ℝ^*K*×*d*^ and ***R*** ∈ ℝ^50×1280^ representation respectively. We project both to a common space ℝ^*p*^ and apply both multiheaded self attention and cross attention to these tensors. Our choice of implementation for these attention modules (MHA) is from ESM-2, utilizing their rotary positional encodings (RoPE). Intuitively this allows codes to be informed by their targets and vice versa. The output of this block are ***E***_*ligand*_ ∈ ℝ^*K*×*p*^, ***E***_*protein*_ ∈ ℝ^50×*p*^ given by the following transformations:

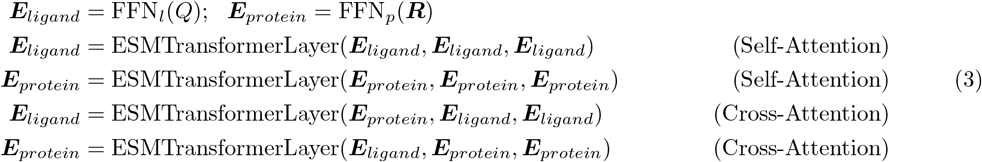

#### Quantized Ligand-Protein Vectorization

After applying self- and cross-attention using ESM-2 Transformer layers, we vectorize the code and protein representations. Codes at this point are ℝ^*K*×*p*^, which we flatten to get 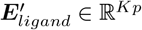. For proteins, across the fixed-length axis (length=50), we compute a softmax-weighted sum to inductively bias towards a (hypothetical) binding site. Finally we project each representation to a shared ℝ^*p*^ space with ReLU activation, and compute the probability of interaction as their cosine similarity (**Figure 1E**).

#### Datasets and Training

The rapid progress in DTI research has produced multiple valuable datasets, but with distinct data imbalances and coverage. These datasets also remain largely disconnected, complicating systematic evaluation of DTI methods. We addressed this by creating MooDengDB, a comprehensive database (named after a pleasantly plump pygmy hippo) that combines and curates data from PLINDER [8], BindingDB [24], BIOSNAP [26], Davis [6], and LCIdb [14]. We are releasing MooDengDB, its training, validation, and test splits, and also a second test set containing out-of-distribution ligands to facilitate reproducible benchmarking across the field (**Table 1**).

**Table 1:**
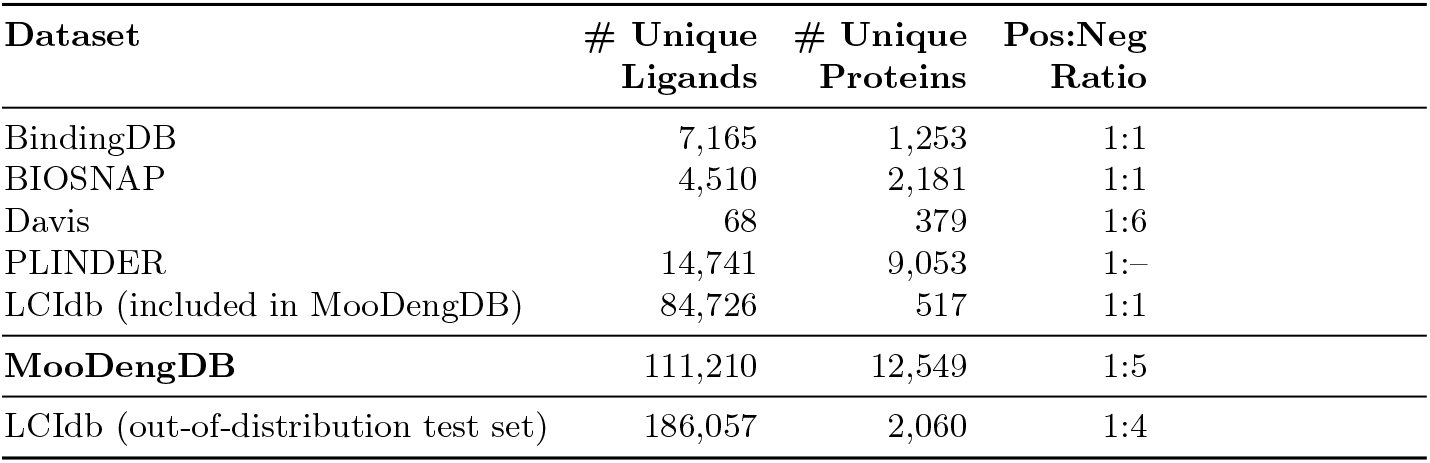
Statistics of Drug-Target Interaction (DTI) datasets.

| Dataset | # Unique<br>Ligands | # Unique<br>Proteins | Pos:Neg<br>Ratio |
| --- | --- | --- | --- |
| BindingDB | 7,165 | 1,253 | 1:1 |
| BIOSNAP | 4,510 | 2,181 | 1:1 |
| Davis | 68 | 379 | 1:6 |
| PLINDER | 14,741 | 9,053 | 1:– |
| LCIdb (included in MooDengDB) | 84,726 | 517 | 1:1 |
| **MooDengDB** | 111,210 | 12,549 | 1:5 |
| LCIdb (out-of-distribution test set) | 186,057 | 2,060 | 1:4 |

Our curation prioritized three key objectives. First, to ensure rigorous evaluation of protein generalizability, we used MMSeqs2 [34] to create training, validation, and test splits (80:10:10) with less than 50% sequence identity between proteins across splits. Second, we addressed the varying data imbalances across datasets. For instance, PLINDER lacked negative examples while LCIdb has a very low protein-to-ligand ratio (1:135). Third, we sought a stringent evaluation of the generalizability of our codebook approach. A good codebook-based prediction should perform well on the challenging out-of-distribution (OOD) case where not only is the test ligand unseen in the training data but it also belongs to a code that itself had no training examples. However, creating an OOD test set presents a circularity: identifying which codes have no training data is impossible until the model is trained and the codes are learned. To address this, we held out a very large number (186,454) of ligands from LCIdb, more than are actually present in MooDengDB. This offers two advantages: it addresses LCIdb’s relative protein-to-ligand imbalance against other datasets when constructing MooDengDB, and makes it highly probable that at least some held-out ligands will be OOD, enabling a robust evaluation of our approach. Finally, we augmented our negative examples with random DTI pairs to obtain a 1:5 positive-to-negative ratio, in keeping with some of the previous work.

CoNCISE was trained using PyTorch’s AdamW optimizer, learning rate of 3 × 10^−4^, and batch size of 32. CoNCISE was trained for 20 epochs across all datasets. We selected the epoch with the highest validation AUPR, waiting at least 10 epochs to ensure broad code coverage.

## 3 Results

### CoNCISE performs state-of-the-art DTI prediction

We evaluated CoNCISE against ConPLex [32], based on PLM-based contrastive learning, and Komet [14], which uses Kronecker factorization of paired protein-ligand features. To our knowledge, these are currently the best-performing sequence-based approaches. In fact, Luo et al reported that ConPLex outperformed many structure-based methods on kinase-drug interaction prediction [25].

Our principal evaluation is on our curated MooDengDB, which combines multiple datasets. To contextualize our results against previous work which used smaller databases, we separately also trained and tested on BioSNAP and BindingDB, using similar train-val-test splitting procedures as for MooDengDB. We did not perform separate evaluations on Davis (too small), PLINDER (lacks negative examples in the original data), and LCIdb (much of it held out for for out-of-distribution testing); all of these were part of MooDengDB. Additionally, to test whether unsupervised clustering of ligands could match the binding-informed codebook, we included a baseline that applies k-means (*k*=4096) clustering to Morgan fingerprints.

As shown in **Table 2**, the results reveal an interesting pattern. On the smaller datasets BioSNAP (32K DTIs) and BindingDB (27K DTIs), CoNCISE performs comparably to existing methods, likely because limited data constrains the learning of its attention-based cross-modal interactions. However, on the substantially larger MooDengDB (766K DTIs), CoNCISE achieves a significant improvement of approximately 15% (AUPR 0.691 vs 0.601 for ConPLex, the second-best method). The k-means baseline performed substantially worse (AUPR 0.435), confirming that binding-informed codebook learning is crucial.

**Table 2:**
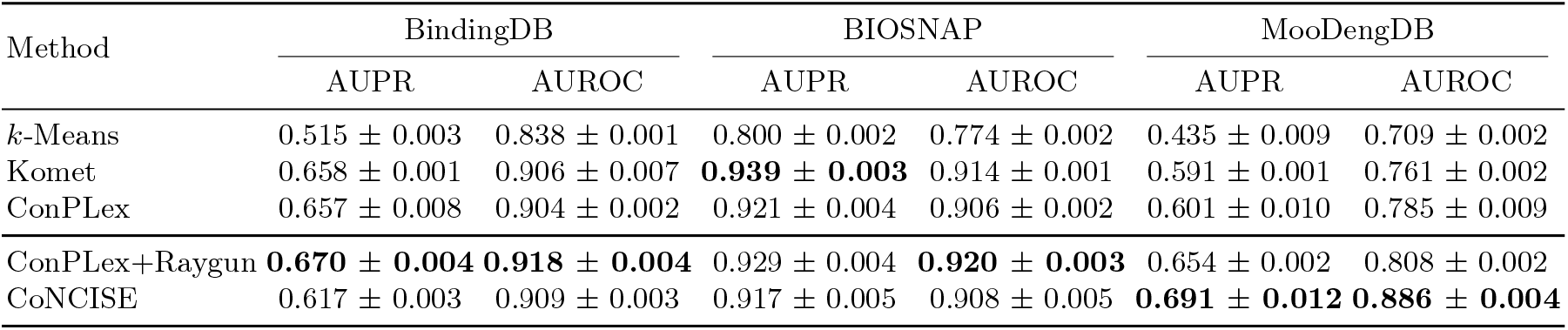
Summary of AUPR and AUROC across different methods and datasets.

To disentangle the contributions of our innovations, we created a “ConPLex-Raygun” variant by modifying ConPLex to use Raygun’s fixed-length protein representations. While this improved upon vanilla ConPLex, it still significantly underperformed CoNCISE on MooDengDB, suggesting that both our hierarchical ligand quantization and the Raygun integration contribute to ConCISE’s gains.

### Ablations for CoNCISE

Given our target space of 32, 768 codes, a key architectural question is how to distribute these codes across layers. Wide architectures (using fewer, wider layers) offer parameter efficiency but sacrifice hierarchical organization, while deep architectures (many narrow layers) enable fine-grained clustering but risk arbitrary subdivisions. We investigated this tradeoff on MooDengDB by comparing three approaches (**Table 3**): a single-layer model producing all 2^15^ codes directly, a 15-layer binary tree model, and CoNCISE’s balanced 3-tier approach. While the deep model (AUPR 0.675) outperformed the flat model (AUPR 0.641), CoNCISE’s balanced architecture proved most effective. This aligns with Mentzer et al. [27]’s observation that FSQ architectures degrade with increased codebook width. We further validated our architecture through a narrow model (1,000 codes) compared against *k*-means clustering. While not as performant as the larger codebooks (AUPR 0.622), it still significantly outperformed *k*-means (AUPR 0.435 using 4,096 clusters; Table 2), demonstrating the crucial value of binding-informed clustering.

**Table 3:** Ablation study of various codebook sizes and topologies.

| Scheme | Flat |  | Hierarchical |  |
| --- | --- | --- | --- | --- |
|  | Wide | Narrow | Deep | CoNCISE |
| Topology | [1 layer] | [1 layer] | [15-layer tree, fan-out=2] | [3-layer tree, fan-out=32] |
| Codebook size | 32,768 | 1,000 | 32,768 | 32,768 |
| AUPR | 0.641 ± 0.018 | 0.622 ± 0.008 | 0.675 ± 0.008 | **0.691 ± 0.012** |
| AUROC | 0.845 ± 0.006 | 0.833 ± 0.004 | 0.857 ± 0.003 | **0.886 ± 0.004** |

### Exploring the CoNCISE representation of small-molecule drug space

CoNCISE organizes ligands into a hierarchical tree structure, where each node represents a unique code combination. Child nodes (e.g., *a*.*b*.*c*) form proper subsets of their parent nodes (e.g., *a*.*b*) (**Figure 2A**). Our first validation confirms that this representation avoids “code collapse”– each FSQ block fully utilizes its 32 segments (**Figure 2B**).

Even though CoNCISE is supervised solely on drug-target binding, we expect it to implicitly capture structural similarities between ligands. This relationship between binding and structural similarity motivated our design– while structurally similar molecules often share binding properties, this correlation is imperfect. Thus, while structural similarity should be an *emergent* property of the model, the underperformance of k-means clustering in DTI prediction highlights that structural similarity alone is insufficient (**Table 2**). We hypothesized that ligands sharing more specific codes (lower in the hierarchy) would show greater structural similarity. We tested this systematically using two experiments. We used Tanimoto scores as our structural similarity metric and restricted our analysis to MooDengDB ligands (**Appendix 4**). The first experiment examined structural similarity between sibling nodes at each tier. We computed pairwise Tanimoto-based similarity scores between siblings, generating 32 × 32 heatmaps at each level. For L1, siblings were the 32 top-level codes; for L2 and L3, we demonstrate on siblings under the “18.*.*” and “18.31.*” subtrees respectively (**Figure 2C**). The average Tanimoto scores between siblings increased with depth, confirming that our representations capture meaningful structural hierarchies.

The second experiment compared how much within-group structural similarity increased from parent to child nodes. We quantified this using the log-ratio of within-group Tanimoto scores— positive values indicate children have greater structural similarity than their parents. The resulting heatmaps showed significant enrichment in lower tiers: 71% of L1-to-L2 comparisons and 63% of L2-to-L3 comparisons had positive log-ratios. As an example, we demonstrate this increasing structural similarity through the “18.*.*” subtree.

### CoNCISE unlocks proteome-scale DTI inference

By compressing the vast ligand space into discrete codes, CoNCISE enables comprehensive protein-ligand binding analysis at an unprecedented scale. We demonstrate this by analyzing the entire human proteome (19,880 proteins from SwissProt after some filtering), computing binding confidence scores for all protein-code combinations. <u>To our knowledge, this is the first comprehensive chemical space-scale analysis of the small-molecule binding preferences of human proteins</u>. This massive computation—approximately 650 million predictions—completed in just 5 hours.

Using a confidence threshold of ≥ 0.95 for predicting binding, we found that most proteins can bind many ligand codes: the 25th to 75th percentiles span 30 to 392 codes (median: 144) (**Figure 3A**). Intriguingly, some proteins (1552) showed no strong binding affinity to any code. The list of these proteins, which may correspond to “undruggable” proteins discussed in the field, is provided in Appendix C. Notably, this number increases when considering only codes that contain some data in MooDengDB’s training set (111,150 ligands), suggesting that some proteins currently considered undruggable may have binding partners in unexplored regions of chemical space. This lends support to our motivation that access to larger compound databases could make it easier to identify drugs for hitherto difficult-to-drug targets.

**Fig. 3:**
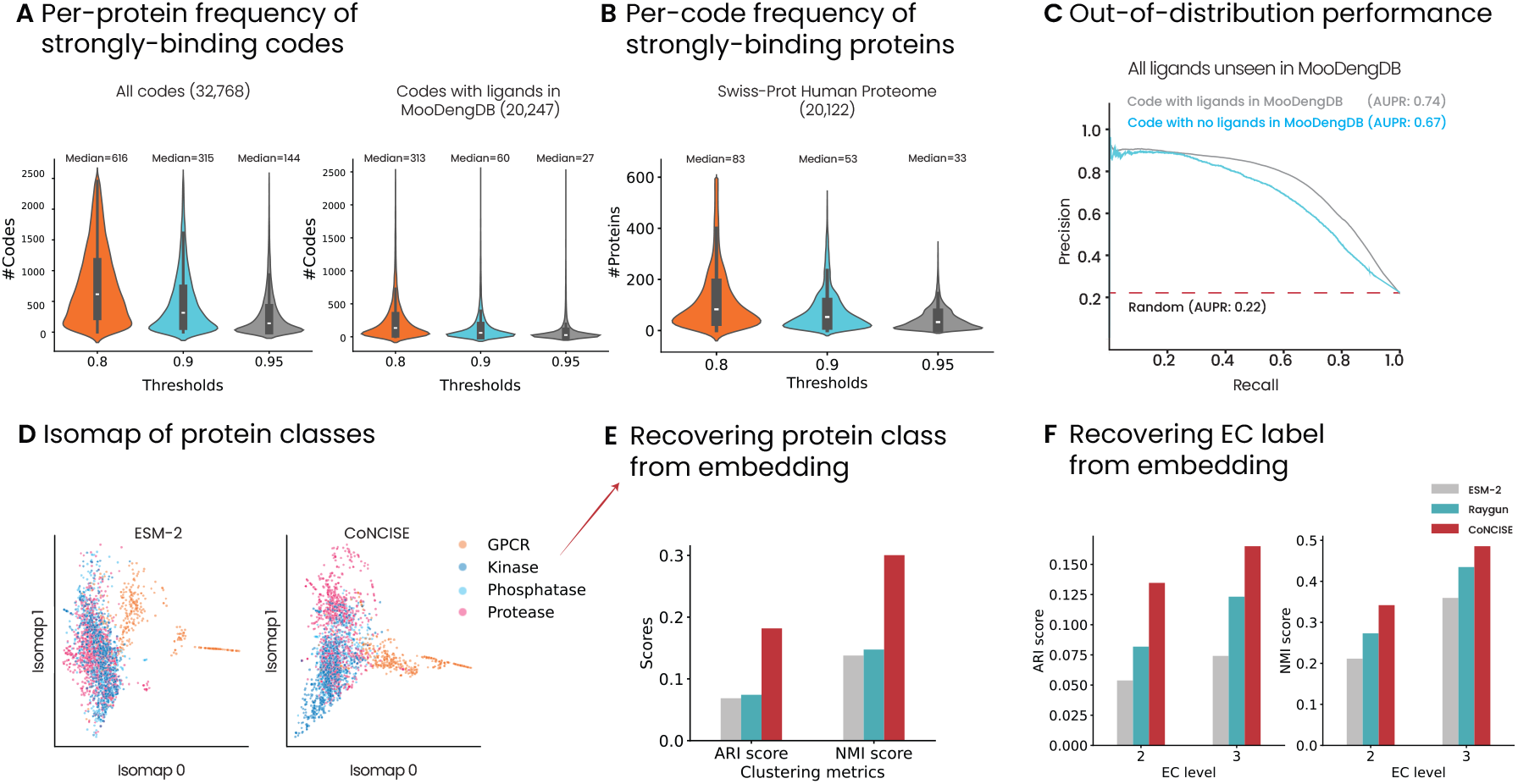
CoNCISE enables proteome-scale analysis of drug binding. A) Most proteins bind multiple codes strongly (median: 144), with some showing no strong binders while others bind >1,500 codes. B) The median code binds 33 human proteins strongly, enabling estimation of potential off-target effects. C) CoNCISE maintains high accuracy on out-of-distribution ligands, whether they map to observed codes (AUPR 0.74) or entirely new codes (AUPR 0.67). D) CoNCISE embeddings provide better separation of key protein classes than ESM-2. E,F) Quantitative validation shows that CoNCISE’s protein representations better capture functional similarities, measured through protein class clustering (E) and enzyme classification hierarchy (F). ARI (adjusted Rand index) and NMI (normalized mutual information) measure clustering overlap with ground-truth class or EC labels.

At the other extreme, we identified 401 highly promiscuous proteins that bind more than 1,500 different codes. Enrichment analysis of these proteins (**Appendix Figure C.1**) revealed significant overlap with metabolic pathways and enzymatic activities. Notably, one enriched group was cytochrome p450, a family of enzymes well-known for their promiscuous binding [15,41], further illustrating CoNCISE’s ability to capture meaningful binding patterns.

We also analyzed the distribution from the perspective of codes, finding that each code typically shows high binding affinity (≥ 0.95 confidence) to approximately 33 human proteins (**Figure 3B**). This finding also has direct implications for drug-discovery: the consideration of off-target effects of a drug should first focus on other proteins that also bind tightly to its code. Conversely, the frequent occurrence of codes binding multiple proteins suggests an intriguing application: developing multi-target therapeutics by identifying and sampling ligands from codes with high predicted affinity to several desired targets simultaneously.

### The CoNCISE representation performs well on diverse, unseen drug categories

The vast chemical space of potential drugs suggests that current experiments sample only a subset of possible DTIs. Thus, training CoNCISE on even a database of substantial size, like MooDengDB, will likely populate only a subset of CoNCISE’s possible 2^15^ codes. This is by design— it allows the model to extend effectively to larger databases like ZINC and Enamine, which likely contain some ligands with remarkably different chemical and binding properties. To systematically evaluate if CoNCISE’s DTI prediction accuracy extends to these unseen drug categories, we had created a challenging out-of-distribution (OOD) test set from LCIdb that only contains ligands excluded from MooDengDB (**Table 1**). Here we analyze CoNCISE performance on this OOD data by further dividing it into two categories: Code-Seen, containing new ligands that map to codes observed during training, and Code-Unseen, containing ligands in previously unobserved codes.

The results, shown in **Figure 3C**, validate CoNCISE’s extensibility in both scenarios. First, the AUPR for the Code-Seen dataset was 0.74, in fact higher than seen on the MooDengDB test set (AUPR 0.69, Table 2), and confirming that the observed codes can be reliably expanded to incorporate new ligands. Second, even the performance on the Code-Unseen dataset (AUPR 0.67), while less strong, was still comparable to the performance on the MooDengDB test set and far higher than random (AUPR 0.22). This robust out-of-distribution performance suggests we successfully achieved our key design goal that CoNCISE’s hierarchical representation should effectively scale to diverse new drug categories while maintaining high DTI accuracy.

### CoNCISE learns functionally-informative protein representations

CoNCISE learns a shared representation space for both drugs and proteins, enabling its protein embeddings to capture binding-relevant features. This motivation is compelling as Raygun, which forms the basis of CoNCISE’s protein representation module, was already shown to better capture protein function than ESM-2 [7]. We therefore investigated whether CoNCISE’s training with drug information could enhance this capability further.

We evaluated this hypothesis through two complementary analyses. First, we examined four protein classes of particular relevance to drug discovery—kinases, phosphatases, GPCRs, and proteases. Isomap [37] visualizations (**Figure 3D**) show that CoNCISE embeddings provide clearer separation between these classes than ESM-2. Quantitatively, we tested this by using each embedding, ESM-2, Raygun, and CoNCISE, to cluster the proteins and assess the agreement of the clustering with ground-truth class labels. As measured by normalized mutual information (NMI) and adjusted Rand index (ARI), CoNCISE demonstrated more than twofold improvement over the other representations (**Figure 3E**).

For our second analysis, we leveraged the Enzyme Commission (EC) classification system, which organizes enzymes in a four-level hierarchy. Since the EC classifications reflect protein binding patterns, they offer an independent validation of CoNCISE’s ability to capture functionally-relevant protein features. Since the top level contains just seven broad categories and the fourth level can be too fine-grained, with very few human proteins per category, we focused on the intermediate levels (2 and 3) which provide useful granularity of enzyme groupings. Again, CoNCISE embeddings showed substantially better clustering by EC number at both levels 2 and 3 (**Figure 3F**). These results demonstrate that CoNCISE’s cross-attention and co-embedding strategy enhances the functional information captured in its already-powerful Raygun protein representations, a crucial capability for accurate DTI prediction.

### CoNCISE enables efficient processing of billion-scale compound databases

The true test of CoNCISE’s scalability lies in its ability to process massive chemical databases like ZINC (600M compounds) and Enamine (6.8B compounds). Despite their size—orders of magnitude larger than MooDengDB—CoNCISE efficiently processed these databases on a single A100 GPU, taking just 3 hours for ZINC and 16 hours for Enamine. We could not compute codes for 5.9% of Enamine (i.e., 0.4B codes) because RDKit failed to compute Morgan fingerprints for them. The resulting 15-bit hierarchical codes occupy less than 10% of the storage space required for the original SMILES representations (**Figure 4C**) and are 0.7% the size of their originating Morgan fingerprints, enabling rapid, memory-efficient virtual screening at unprecedented scale.

**Fig. 4:**
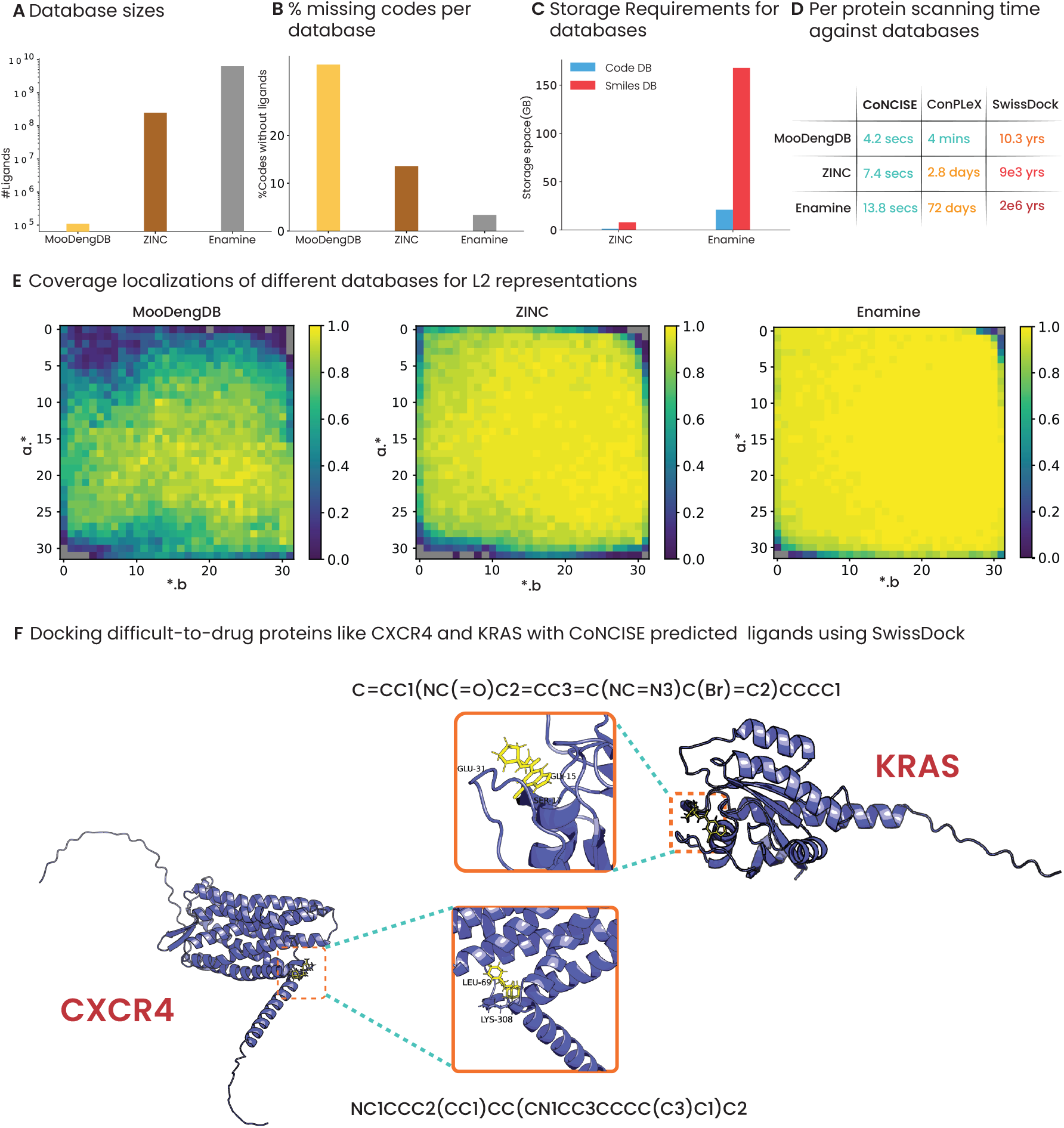
Scaling drug discovery to billion-compound libraries. A-D) CoNCISE enables efficient processing of massive databases: while Enamine contains 6.4B compounds and requires 150GB storage, screening it takes only 13.8 seconds per protein (versus 72 days for ConPLeX). E) Progressive filling of chemical space from MooDengDB to ZINC to Enamine, shown through L2 code coverage. For each L2 code (*a*.*b*), the pixel indicates the fraction of non-empty L3 (*a*.*b*.*\**) codes. F) Proof-of-concept screening of Enamine for difficult-to-drug targets: CoNCISE rapidly identified strong predicted binders for KRAS and CXCR4, which were then filtered through DiffDock and SwissDock.

The coverage analysis reveals how these larger databases progressively fill CoNCISE’s code space. When considering the overall fraction of 32,768 L3 codes that contain at least one ligand, MooDengDB reaches only 65.2% only a fraction of the possible codes, while ZINC is able to cover 86% and Enamine reaches 97% of the total code space. To understand where this improved coverage occurs, we generated 32 × 32 heatmaps showing the occupancy of L3 codes for each L2 representation (**Figure 4E**). These visualizations reveal that MooDengDB’s coverage is sparsest at the boundaries of the chemical space, precisely where ZINC and Enamine contribute many novel compounds.

This comprehensive coverage of chemical space, combined with CoNCISE’s strong performance on out-of-distribution compounds (as shown in the previous section), suggests that our approach can effectively scale to virtual screening of billion-compound libraries while maintaining DTI prediction accuracy. The efficient encoding means that, once processed, these massive databases can be searched in constant time regardless of their size. In fact, the only reason Enamine takes slightly longer to query than MooDengDB is because the size of the output, i.e. the number of ligands matched, is much larger in the former (**Figure 4D**).

### CoNCISE enables efficient virtual screening of billion-scale databases

A key advantage of CoNCISE’s code-based representation is that DTI inference time is effectively decoupled from database size. Once a database is indexed with codes, scanning for high-affinity binders requires only the code representations, enabling near-constant-time virtual screening even across billions of compounds (**Algorithm 5**).

To demonstrate this capability, we conducted virtual screening for three proteins traditionally considered difficult to drug: KRAS, CXCR4, and P53. For each protein, we first identified all codes predicted to bind with high confidence (probability ≥ 0.95), typically yielding around 300 codes per protein. We then randomly selected one Enamine compound per code and screened these candidates using DiffDock [5], followed by SwissDock [4] validation of the highest-scoring ligand and pose (**Figure 4F, D.2**). Notably, the entire process for each protein took less than 2 hours, most of it spent on the multiple DiffDock runs.

The results support CoNCISE’s ability to identify promising candidates efficiently. SwissDock predicted strong binding energies (*Δ*G) for all three proteins (P53: −6.9 kcal/mol, KRAS: −7.2 kcal/mol, CXCR4: −7.2 kcal/mol), corresponding to *K*_*D*_ of approximately 20 *µ*M or better. While these results are promising, we do note some limitations: DiffDock does not account for protein conformational changes between bound and unbound states, and our docking pipeline could be further optimized. However, this proof-of-concept demonstrates CoNCISE’s potential—we examined just one compound per code, and better-directed sampling within promising codes could yield even stronger candidates.

## 4 Discussion

CoNCISE demonstrates that most binding-relevant information in chemical space can be captured in just 15 bits when learned specifically for DTIs. This extreme compression— from the 2^2048^ space of Morgan fingerprints to just 2^15^ codes—– not only massively speeds up inference but also reveals fundamental structure in how small molecules bind with proteins. Our work suggests that while structural similarity between molecules correlates with binding similarity, directly optimizing for structural patterns (e.g., k-means clustering) is less effective than letting these patterns emerge through supervision on binding data.

Our technical innovation of the hierarchical FSQ architecture with residual learning between tiers, could offer broader value beyond DTI prediction in domains where task-relevant hierarchical structure exists in some latent space. Here, the compression enabled the first comprehensive analysis of potential small-molecule binding partners across the human proteome, offering new perspectives on protein druggability. Our analysis suggests that some proteins considered “undruggable” may simply have binding partners in unexplored regions of chemical space, highlighting the value of efficient search through massive compound libraries.

CoNCISE’s compact and interpretable organization of the small-molecule space enables several practical advances in drug discovery and cellular biology. Most directly, it democratizes access to large-scale virtual screening—researchers can now query massive compound libraries like Enamine (6.4B molecules) in seconds rather than weeks. Beyond that, our approach enables new experimental design strategies. For instance, CoNCISE could enable the parsimonious design of drug panels: rather than testing compounds randomly, one could systematically sample from different code groups to maximize coverage of binding patterns.

The power of our approach could be further enhanced in several directions. While we focused on sequence-based prediction for its speed and applicability, incorporating additional structural and mechanistic information could improve performance. Additionally, while we demonstrated utility in drug discovery, similar approaches could guide the design of targeted perturbations for studying cellular processes. For example, when seeking to perturb a pathway of interest but being unsure where perturbation would be most effective, CoNCISE could help researchers identify a small set of compounds predicted to collectively modulate all proteins in a pathway of interest. Moreover, CoNCISE’s codebook-based approach provides a framework for incorporating functional considerations such as inhibition and pharmacokinetics– just as our codes learned to capture binding patterns, they could be fine-tuned to reflect these additional pharmacological properties.

Looking forward, CoNCISE opens exciting possibilities for systematic design of chemical tools in biology, from multi-target therapeutics to pathway-specific perturbation panels. By making large-scale virtual screening both accessible and accurate, it represents a significant step toward democratizing drug discovery.

## Code and Data availability

The CoNCISE code and the MooDengDB is publicly available at https://github.com/rohitsinghlab/ConCISE.

## Acknowledgements

We thank the Tufts BCB group for helpful comments. L.V. thanks the DIAMONDS REU program at Tufts (NSF grant 2149871).

## A Model Architecture

The CoNCISE architecture comprises two major components: a) the drug encoder composed of FSQ blocks, and b) DTI layer, that uses the transformed drug and protein embeddings to predict binding affinities. We describe these two layers in detail below:

### A.1 Drug Encoder

Excluding parameter-free FSQ layers, the drug encoder module has two main hyper-parameters: the embedding dimension *d* and the number of FSQ blocks *K*. As the first step in the forward process, the drug encoder linearly projects the 2,048-dimensional fingerprint vector into a much smaller *d*−dimensional space. The projected result then passes through the *K* FSQ blocks, which sandwiches a parameter-free FSQ layer between linear down-sampling and up-sampling layers. The downsampling dimension is dependent on the choice of the parameter-free FSQ layers’ hyper-parameters. The forward operations of an individual *FSQ* block is shown in **Algorithm 1**.

#### Algorithm 1: Forward Pass of FSQBlock

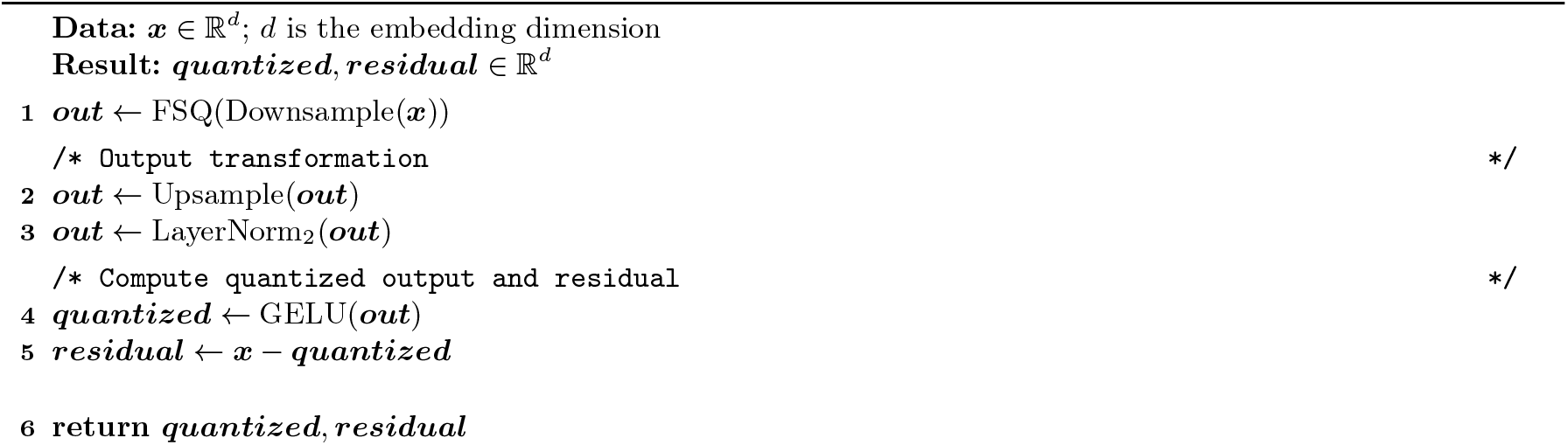

In our final design, we set *K* = 3 and *d* = 256. The forward operation of complete drug encoder module comprising 3 FSQ blocks is shown in **Algorithm 2**.

#### Algorithm 2: Forward Pass of Residual Quantization Model (Drug Encoder)

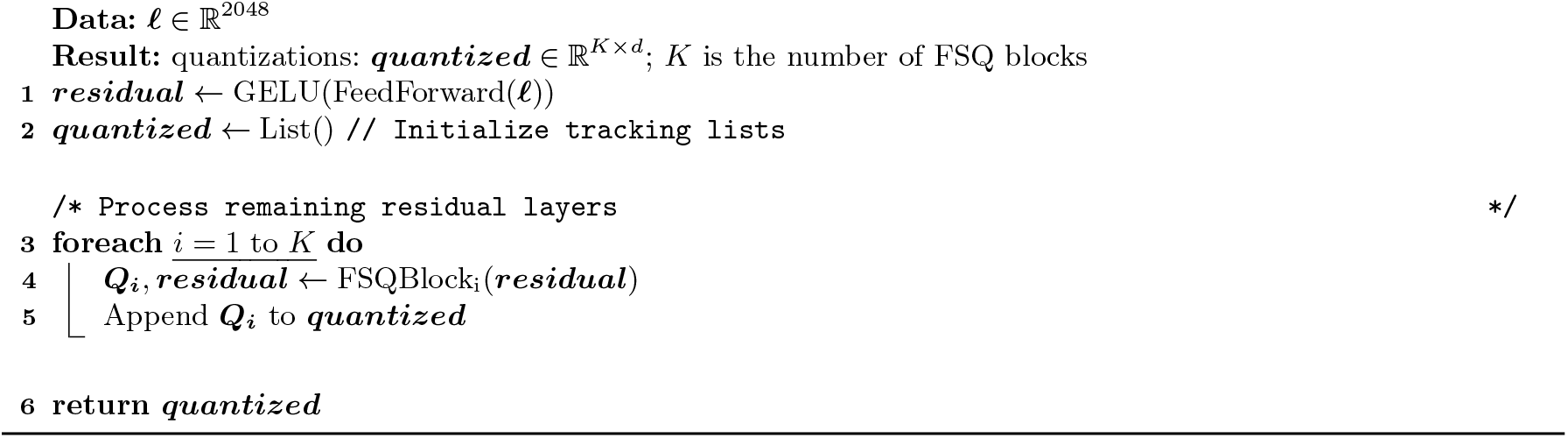

### A.2 DTI module

A significant portion of the learnable parameters in the CoNCISE DTI module is concentrated in its self- and cross-attention layers, which are designed to capture binding-specific relationships between protein and drug embeddings. Initially, the DTI module passes the outputs of the drug and protein encoders through two self-attention layers to capture intra-embedding associations within each type. Then, the two cross-attention layers facilitate information exchange between the drug and protein embeddings. Finally, the cross-attention outputs are condensed into a 1-dimensional representation, and their normalized inner product is returned as the predicted affinity probability. The forward operation of the DTI module is described in Algorithm 3.

#### Algorithm 3: Forward Pass of Binding Prediction Model

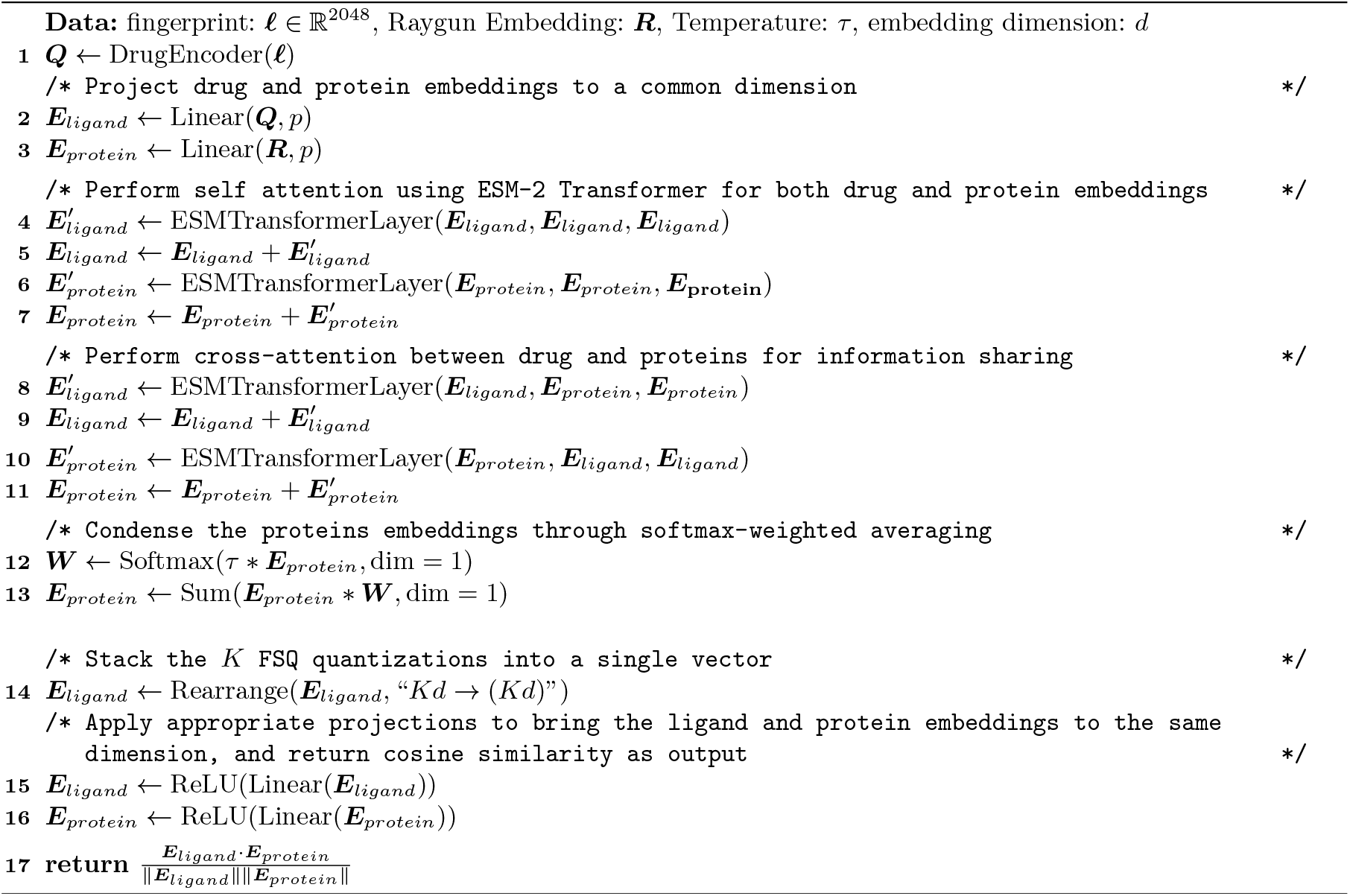

### A.3 Converting L3 code representations back to the quantized code embeddings

The original implementation of the parameter-free FSQ layer gave users the functionality to input integer codes representing possible quantizations and obtain the resulting quantized embeddings as output. This feature is a simple mapping that goes between an integer index and a point on the grid line. We used this feature to obtain the code embeddings.

Given a L3 code representation “*a*_1_.*a*_2_.*a*_3_”, we used the code *a*_*i*_ to obtain the FSQ_*i*_’s quantized embeddings, passing it through the upsampling projectors immediately after the FSQ layers. These individual embeddings were then concatenated to obtain the corresponding drug projector embedding in a completely loss-less manner.

## B Measuring structural homology between ligand representations

We used Tanimoto score to compute chemical similarities between small molecules. Given extended connectivity morgan fingerprint (ECFP4) representations for two smiles: *F*_1_, *F*_2_ ∈ {0, 1} ^2048^, the Tanimoto score is simply the Jaccard similarity computed between binary vectors. In other words,

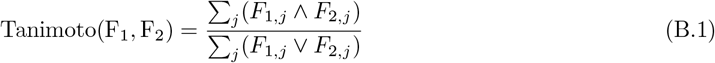

Tanimoto score, however, is only limited to finding similarities between small molecules. We extended this metric further to obtain a statistical measure of similarities between two ligand groups. Let *L*_1_ and *L*_2_ be the set of ligands represented by groups associated with our discretized code representations. Then, the algorithm for computing the group similarity is described below:

### Algorithm 4: Algorithm to measure code similarities

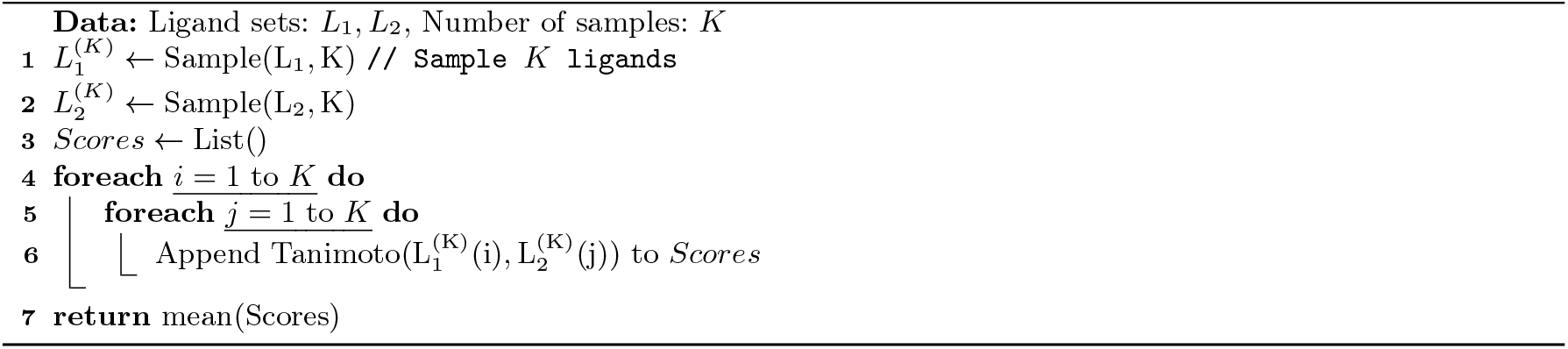

### B.1 Log-ratio of Tanimoto similarities

We compared the structural enrichment between the parent and child nodes by computing their Tanimoto similarities, using the process described in **Algorithm 4** and computing the log ratio:

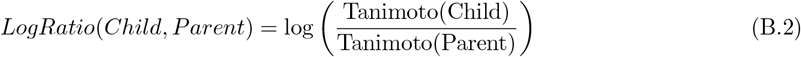

A value greater than 0 implies that the ligands in the child nodes are more structurally enriched than their parents.

## C Undruggable and Promiscuous proteins

We provide the link to the 1552 SwissProt human proteins that CoNCISE found to lack high affinity binding codes (when threshold is set to 0.95) in our github url: https://github.com/rohitsinghlab/CoNCISE/blob/main/data/predicted-undruggables.txt

The 401 promiscuous codes are also provided in the github url: https://github.com/rohitsinghlab/CoNCISE/blob/main/data/predicted-promiscuous.tsv. We additionally performed enrichment analysis on these 401 proteins, the results of which are shown in **Figure C.1**. As expected, these protein candidates

**Fig. C.1:**
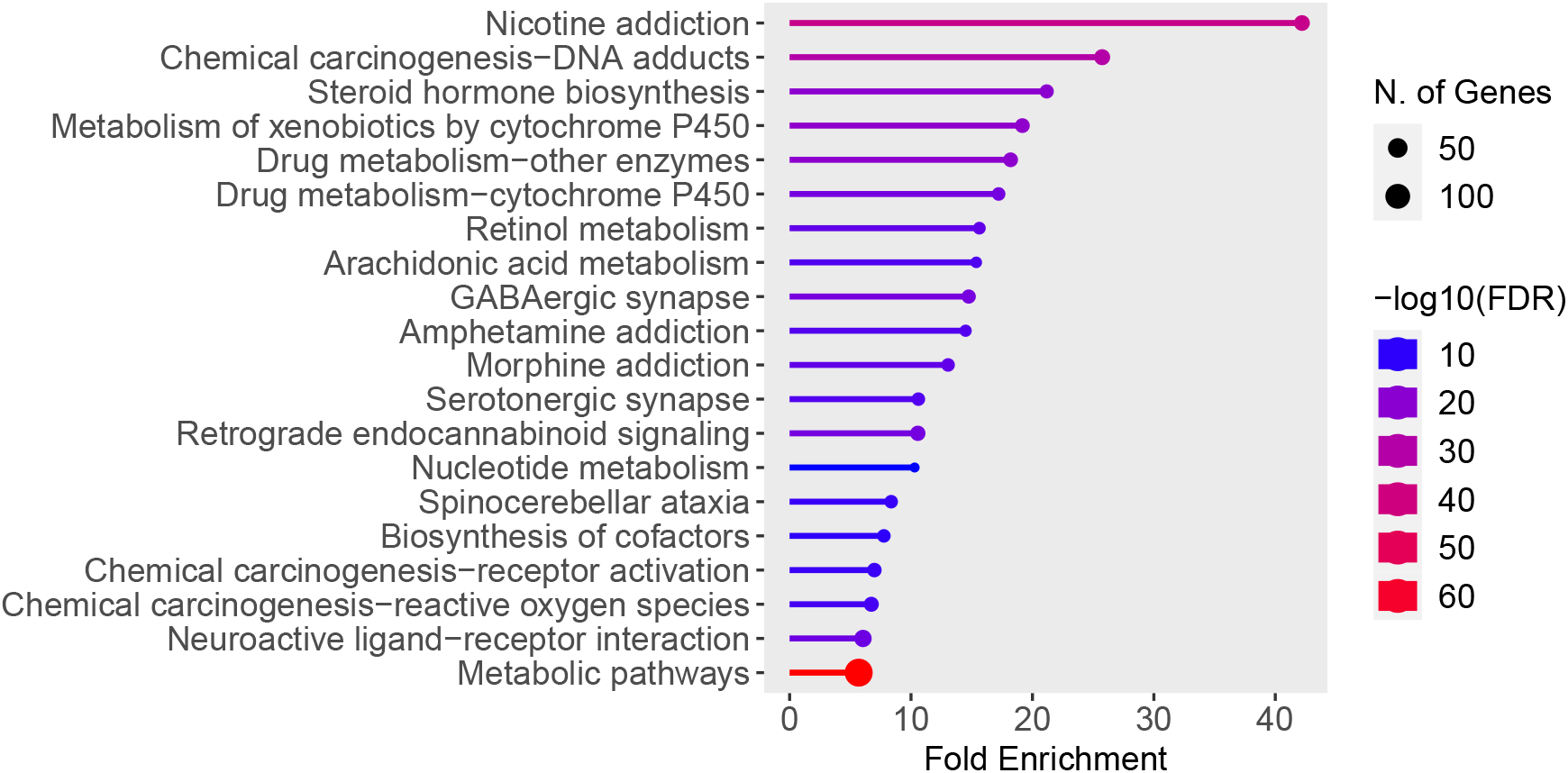
Enrichment results for the 401 promiscuous proteins identified by CoNCISE. Figure obtained from ShinyGO 0.80 [11]

## D Database-wide ligand screening to find high affinity binding targets

After the construction of ligand-code database for Enamine, the process of finding binding candidates can be decoupled into two separate steps: a) finding codes that have a high binding affinity to the protein, and b) searching through the ligand-code database to find ligands mapped to the high affinity codes. The overall process is described in Algorithm 5. Since the process of finding high affinity codes, given a protein, can essentially be done in a constant time, and the database-lookup step is roughly constant-time, the overall screening process for a protein can be done very fast.

### Algorithm 5: Algorithm to scan for high-affinity binders from a ligand database

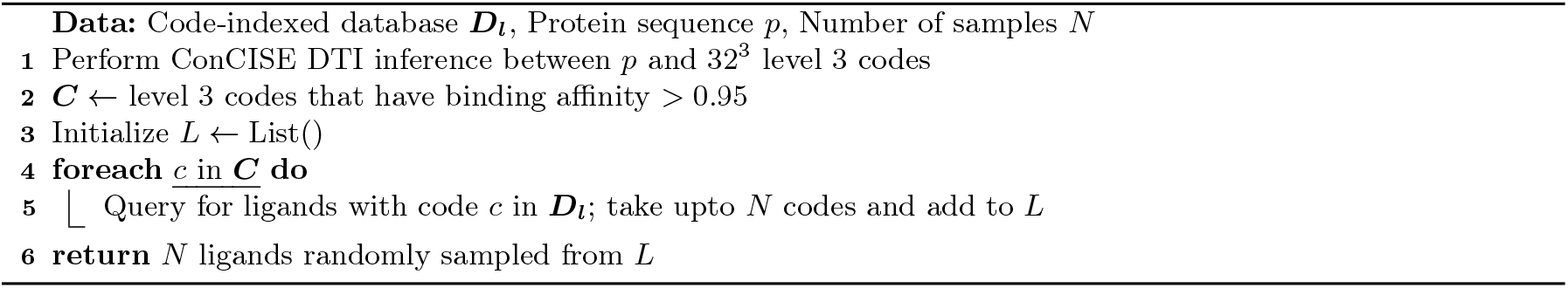

**Fig. D.2:**
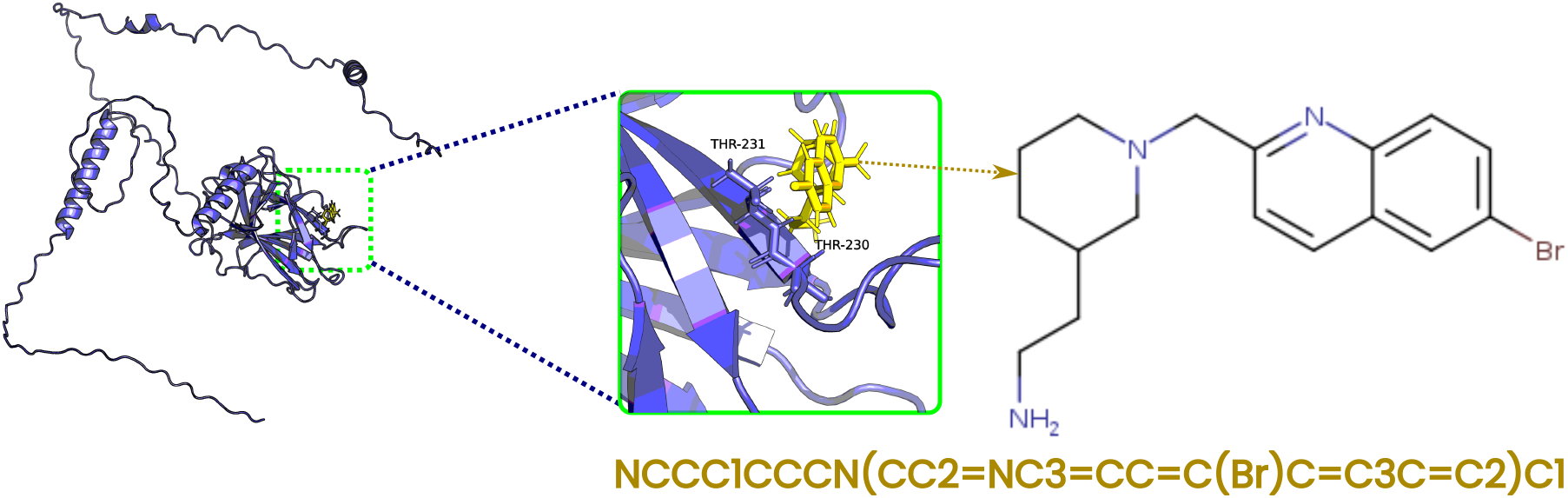
Docking monomeric P53 with a CoNCISE-predicted ligand binder using SwissDock

