## Supplementary document for "Learning a CoNCISE language for small-molecule binding"

### A Model Architecture

The CoNCISE architecture comprises two major components: a) the drug encoder composed of FSQ blocks, and b) DTI layer, that uses the transformed drug and protein embeddings to predict binding affinities. We describe these two layers in detail below:

#### A.1 Drug Encoder

Excluding parameter-free FSQ layers, the drug encoder module has two main hyper-parameters: the embedding dimension $d$ and the number of FSQ blocks $K$. As the first step in the forward process, the drug encoder linearly projects the 2,048-dimensional fingerprint vector into a much smaller $d-$dimensional space. The projected result then passes through the $K$ FSQ blocks, which sandwiches a parameter-free FSQ layer between linear down-sampling and up-sampling layers. The downsampling dimension is dependent on the choice of the parameter-free FSQ layers' hyper-parameters. The forward operations of an individual $FSQ$ block is shown in **Algorithm** 1.

**Algorithm 1:** Forward Pass of FSQBlock

**Data:** $\boldsymbol{x} \in \mathbb{R}^d$; $d$ is the embedding dimension
**Result:** $\boldsymbol{quantized}, \boldsymbol{residual} \in \mathbb{R}^d$

$\boldsymbol{out} \leftarrow \text{FSQ}(\text{Downsample}(\boldsymbol{x}))$
`/* Output transformation */`
$\boldsymbol{out} \leftarrow \text{Upsample}(\boldsymbol{out})$
$\boldsymbol{out} \leftarrow \text{LayerNorm}_2(\boldsymbol{out})$
`/* Compute quantized output and residual */`
$\boldsymbol{quantized} \leftarrow \text{GELU}(\boldsymbol{out})$
$\boldsymbol{residual} \leftarrow \boldsymbol{x} - \boldsymbol{quantized}$

**return** $\boldsymbol{quantized}, \boldsymbol{residual}$

In our final design, we set $K = 3$ and $d = 256$. The forward operation of complete drug encoder module comprising 3 FSQ blocks is shown in **Algorithm** 2.

**Algorithm 2:** Forward Pass of Residual Quantization Model (Drug Encoder)

**Data:** $\boldsymbol{\ell} \in \mathbb{R}^{2048}$
**Result:** quantizations: $\boldsymbol{quantized} \in \mathbb{R}^{K \times d}$; $K$ is the number of FSQ blocks

$\boldsymbol{residual} \leftarrow \text{GELU}(\text{FeedForward}(\boldsymbol{\ell}))$
$\boldsymbol{quantized} \leftarrow \text{List}()$ `// Initialize tracking lists`

`/* Process remaining residual layers */`
**foreach** $i = 1$ to $K$ **do**
  $\boldsymbol{Q_i}, \boldsymbol{residual} \leftarrow \text{FSQBlock}_\text{i}(\boldsymbol{residual})$
  Append $\boldsymbol{Q_i}$ to $\boldsymbol{quantized}$

**return** $\boldsymbol{quantized}$

#### A.2 DTI module

A significant portion of the learnable parameters in the CoNCISE DTI module is concentrated in its self- and cross-attention layers, which are designed to capture binding-specific relationships between protein and drug embeddings. Initially, the DTI module passes the outputs of the drug and protein encoders through two self-attention layers to capture intra-embedding associations within each type. Then, the two cross-attention layers facilitate information exchange between the drug and protein embeddings. Finally, the cross-attention outputs are condensed into a 1-dimensional representation, and their normalized inner product is returned as the predicted affinity probability. The forward operation of the DTI module is described in Algorithm 3.

**Algorithm 3:** Forward Pass of Binding Prediction Model

**Data:** fingerprint: $\boldsymbol{\ell} \in \mathbb{R}^{2048}$, Raygun Embedding: $\boldsymbol{R}$, Temperature: $\tau$, embedding dimension: $d$

$\boldsymbol{Q} \leftarrow \text{DrugEncoder}(\boldsymbol{\ell})$

`/* Project drug and protein embeddings to a common dimension */`

$\boldsymbol{E}_{ligand} \leftarrow \text{Linear}(\boldsymbol{Q}, p)$

$\boldsymbol{E}_{protein} \leftarrow \text{Linear}(\boldsymbol{R}, p)$

`/* Perform self attention using ESM-2 Transformer for both drug and protein embeddings */`

$\boldsymbol{E}'_{ligand} \leftarrow \text{ESMTransformerLayer}(\boldsymbol{E}_{ligand}, \boldsymbol{E}_{ligand}, \boldsymbol{E}_{ligand})$

$\boldsymbol{E}_{ligand} \leftarrow \boldsymbol{E}_{ligand} + \boldsymbol{E}'_{ligand}$

$\boldsymbol{E}'_{protein} \leftarrow \text{ESMTransformerLayer}(\boldsymbol{E}_{protein}, \boldsymbol{E}_{protein}, \boldsymbol{E}_{\mathbf{protein}})$

$\boldsymbol{E}_{protein} \leftarrow \boldsymbol{E}_{protein} + \boldsymbol{E}'_{protein}$

`/* Perform cross-attention between drug and proteins for information sharing */`

$\boldsymbol{E}'_{ligand} \leftarrow \text{ESMTransformerLayer}(\boldsymbol{E}_{ligand}, \boldsymbol{E}_{protein}, \boldsymbol{E}_{protein})$

$\boldsymbol{E}_{ligand} \leftarrow \boldsymbol{E}_{ligand} + \boldsymbol{E}'_{ligand}$

$\boldsymbol{E}'_{protein} \leftarrow \text{ESMTransformerLayer}(\boldsymbol{E}_{protein}, \boldsymbol{E}_{ligand}, \boldsymbol{E}_{ligand})$

$\boldsymbol{E}_{protein} \leftarrow \boldsymbol{E}_{protein} + \boldsymbol{E}'_{protein}$

`/* Condense the proteins embeddings through softmax-weighted averaging */`

$\boldsymbol{W} \leftarrow \text{Softmax}(\tau * \boldsymbol{E}_{protein}, \text{dim} = 1)$

$\boldsymbol{E}_{protein} \leftarrow \text{Sum}(\boldsymbol{E}_{protein} * \boldsymbol{W}, \text{dim} = 1)$

`/* Stack the` $K$ `FSQ quantizations into a single vector */`

$\boldsymbol{E}_{ligand} \leftarrow \text{Rearrange}(\boldsymbol{E}_{ligand}, \text{“}Kd \rightarrow (Kd)\text{”})$

`/* Apply appropriate projections to bring the ligand and protein embeddings to the same dimension, and return cosine similarity as output */`

$\boldsymbol{E}_{ligand} \leftarrow \text{ReLU}(\text{Linear}(\boldsymbol{E}_{ligand}))$

$\boldsymbol{E}_{protein} \leftarrow \text{ReLU}(\text{Linear}(\boldsymbol{E}_{protein}))$

**return** $\frac{\boldsymbol{E}_{ligand} \cdot \boldsymbol{E}_{protein}}{\|\boldsymbol{E}_{ligand}\| \|\boldsymbol{E}_{protein}\|}$

##### A.3 Converting L3 code representations back to the quantized code embeddings

The original implementation of the parameter-free FSQ layer gave users the functionality to input integer codes representing possible quantizations and obtain the resulting quantized embeddings as output. This feature is a simple mapping that goes between an integer index and a point on the grid line. We used this feature to obtain the code embeddings.

Given a L3 code representation "$a_1.a_2.a_3$", we used the code $a_i$ to obtain the $\mathrm{FSQ}_i$'s quantized embeddings, passing it through the upsampling projectors immediately after the FSQ layers. These individual embeddings were then concatenated to obtain the corresponding drug projector embedding in a completely loss-less manner.

#### B Measuring structural homology between ligand representations

We used Tanimoto score to compute chemical similarities between small molecules. Given extended connectivity morgan fingerprint (ECFP4) representations for two smiles: $F_1, F_2 \in \{0,1\}^{2048}$, the Tanimoto score is simply the Jaccard similarity computed between binary vectors. In other words,

$$\mathrm{Tanimoto}(\mathrm{F_1}, \mathrm{F_2}) = \frac{\sum_j (F_{1,j} \wedge F_{2,j})}{\sum_j (F_{1,j} \vee F_{2,j})} \tag{B.1}$$

Tanimoto score, however, is only limited to finding similarities between small molecules. We extended this metric further to obtain a statistical measure of similarities between two ligand groups. Let $L_1$ and $L_2$ be the set of ligands represented by groups associated with our discretized code representations. Then, the algorithm for computing the group similarity is described below:

**Algorithm 4:** Algorithm to measure code similarities

**Data:** Ligand sets: $L_1, L_2$, Number of samples: $K$

$L_1^{(K)} \leftarrow \mathrm{Sample}(\mathrm{L_1}, \mathrm{K})$ `// Sample` $K$ `ligands`
$L_2^{(K)} \leftarrow \mathrm{Sample}(\mathrm{L_2}, \mathrm{K})$
$Scores \leftarrow \mathrm{List}()$
**foreach** $i = 1$ to $K$ **do**
  **foreach** $j = 1$ to $K$ **do**
    Append $\mathrm{Tanimoto}(\mathrm{L_1^{(K)}(i)}, \mathrm{L_2^{(K)}(j)})$ to $Scores$
**return** mean(Scores)

##### B.1 Log-ratio of Tanimoto similarities

We compared the structural enrichment between the parent and child nodes by computing their Tanimoto similarities, using the process described in **Algorithm 4** and computing the log ratio:

$$LogRatio(Child, Parent) = \log\left(\frac{\mathrm{Tanimoto(Child)}}{\mathrm{Tanimoto(Parent)}}\right) \tag{B.2}$$

A value greater than 0 implies that the ligands in the child nodes are more structurally enriched than their parents.

#### C Undruggable and Promiscuous proteins

We provide the link to the 1552 SwissProt human proteins that CoNCISE found to lack high affinity binding codes (when threshold is set to 0.95) in our github url: `https://github.com/rohitsinghlab/CoNCISE/blob/main/data/predicted-undruggables.txt`

The 401 promiscuous codes are also provided in the github url: `https://github.com/rohitsinghlab/CoNCISE/blob/main/data/predicted-promiscuous.tsv`. We additionally performed enrichment analysis on these 401 proteins, the results of which are shown in **Figure C.1**. As expected, these protein candidates showed enrichment for enzymatic and drug activities which are known contain proteins with high promiscuity.

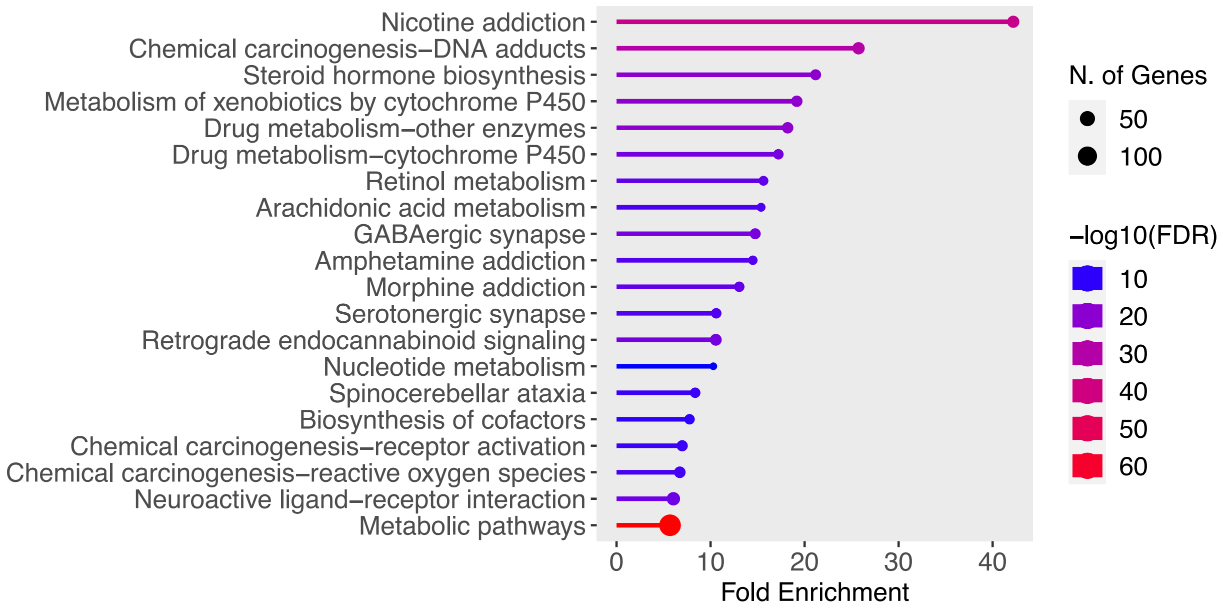


Fig. C.1: Enrichment results for the 401 promiscuous proteins identified by CoNCISE. Figure obtained from ShinyGO 0.80 [11]

### D Database-wide ligand screening to find high affinity binding targets

After the construction of ligand-code database for Enamine, the process of finding binding candidates can be decoupled into two separate steps: a) finding codes that have a high binding affinity to the protein, and b) searching through the ligand-code database to find ligands mapped to the high affinity codes. The overall process is described in Algorithm 5. Since the process of finding high affinity codes, given a protein, can essentially be done in a constant time, and the database-lookup step is roughly constant-time, the overall screening process for a protein can be done very fast.

**Algorithm 5:** Algorithm to scan for high-affinity binders from a ligand database

**Data:** Code-indexed database $\boldsymbol{D_l}$, Protein sequence $p$, Number of samples $N$
Perform ConCISE DTI inference between $p$ and $32^3$ level 3 codes
$\boldsymbol{C} \leftarrow$ level 3 codes that have binding affinity $> 0.95$
Initialize $L \leftarrow$ List()
**foreach** $c$ in $\boldsymbol{C}$ **do**
    Query for ligands with code $c$ in $\boldsymbol{D_l}$; take upto $N$ codes and add to $L$
**return** $N$ ligands randomly sampled from $L$

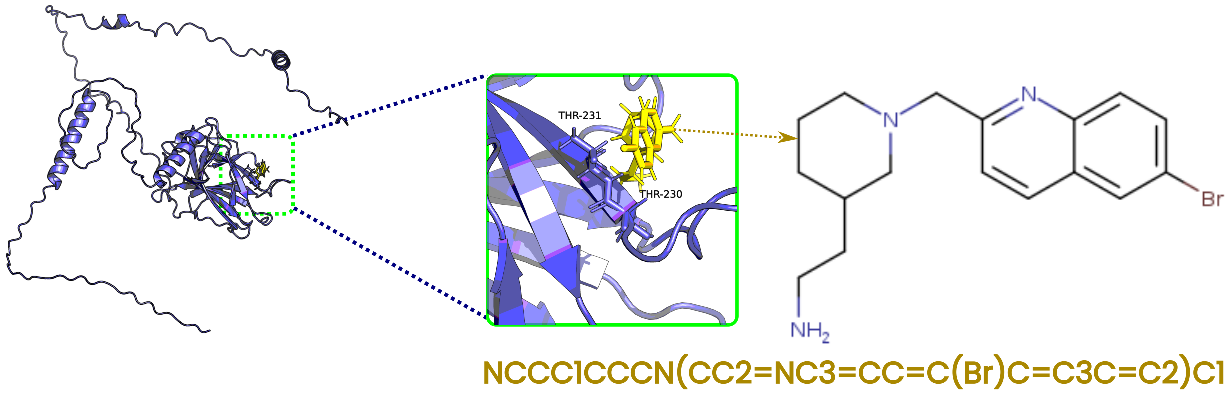


Fig. D.2: Docking monomeric P53 with a CoNCISE-predicted ligand binder using SwissDock
